# Inferring intracellular signal transduction circuitry from molecular perturbation experiments

**DOI:** 10.1101/107730

**Authors:** Michelle L. Wynn, Megan Egbert, Nikita Consul, Jungsoo Chang, Zhi-Fen Wu, Sofia D. Meravjer, Santiago Schnell

## Abstract

The development of network inference methodologies that accurately predict connectivity in dysregulated pathways may enable the rational selection of patient therapies. Accurately inferring an intracellular network from data remains a very challenging problem in molecular systems biology. Living cells integrate extremely robust circuits that exhibit significant heterogeneity, but still respond to external stimuli in predictable ways. This phenomenon allows us to introduce a network inference methodology that integrates measurements of protein activation from perturbation experiments. The methodology relies on logic-based networks to provide a predictive approximation of the transfer of signals in a network. The approach presented was validated *in silico* with a set of test networks and applied to investigate the epidermal growth factor receptor signaling of a breast epithelial cell line, MFC10A. In our analysis, we predict the potential signaling circuitry most likely responsible for the experimental readouts of several proteins in the mitogen activated protein kinase and phosphatidylinositol-3 kinase pathways. The approach can also be used to identify additional necessary perturbation experiments to distinguish between a set of possible candidate networks.

## Introduction

It is increasingly likely that the large number of off-target effects associated with targeted therapeutics is due to the complex interactions and emergent behaviors intrinsic to dysregulated signaling pathways in cells. In order to rationally develop precise therapeutic avenues to target key dysregulated pathways in diseased cells, it is critical to develop methodologies that can help us understand how complex intracellular signaling pathways are wired as an integrated network within both normal and aberrant cells. While targeted molecular inhibitors hold great promise for controlling disease, in the clinic these drugs are often not as successful as their pre-clinical data indicated.

The pathways associated with epidermal growth factor (EGF) receptor signaling are of great clinical interest. EGF receptors can activate both the mitogen activated protein kinase (MAPK) pathway, which promotes motility, invasion, and angiogenic factor production, and the phosphatidylinositol-3 kinase (PI3K) pathway, which controls anchorage independent growth and modulates glucose metabolism (Plas and Thompson 2005; Robey and Hay 2009; van Golen et al. 2002; Zhang et al. 2003). Both the MAPK and PI3K pathways are highly dysregulated in cancer (Halilovic et al. 2010; McCubrey et al. 2006; Meier et al. 2005; Rommel et al. 1999; Won et al. 2012), neuroinflammation (Liu et al. 2014), obesity and type 2 diabetes (Bernal-Mizrachi et al. 2014; Kulkarni et al. 2012; Schultze et al. 2012), and developmental disorders (Hong et al. 2008; Nie and Chang 2007). Therapeutic targeting of the MAPK pathway has been less effective than hoped (McCormick 2011; Won et al. 2012), and several regulatory mechanisms have been suggested to explain the apparent cross-talk between the MAPK and PI3K pathways (Serra et al. 2011). Understanding precisely how these networks are wired in normal and diseased cells will have tremendous clinical significance.

Boolean logic can be used to simulate signal transfer in a molecular and biological network in a manner similar to signal transfer in a digital circuit. Boolean logic models of signaling networks approximate the sigmoidal regulation of a target molecule by an activator or inhibitor (Wynn et al. 2012). They can also be used to explicitly simulate enzyme activation and inhibition in a set of biochemical reactions as well as bistability and chemical hysteresis (Arkin and Ross 1994; Hjelmfelt and Ross 1995; Hjelmfelt et al. 1993). Logic-based models are powerful tools for studying complex systems, such as intracellular molecular networks, because they are qualitatively predictive and do not require parameter information or mechanistic details needed for more quantitatively precise kinetic methods (Albert et al. 2008; Glass and Kauffman 1973; Thomas 2006; Wynn et al. 2012). Dynamic signaling models based on logic networks have high predictive power only if the underlying logic network is accurately constructed. Of course, this is also true of highly quantitative kinetic-based differential equation models, which require knowledge of the underlying circuitry as well as a very large number of unknown parameters. While treating protein activation levels in binary terms of ***ON*** and ***OFF*** is a simplification of the complexity of molecular function within a cell, it is important to emphasize what ***ON*** and ***OFF*** values signify in a logic-based signaling model. Specifically, when a node is in an ***ON*** or ***OFF*** state, the molecule is assumed to be above or below, respectively, the threshold needed to induce an appreciable effect in the molecule or molecules it regulates (Wynn et al. 2012). This threshold assumption is similar to the continuous concentration approach used by Ross and colleagues (Arkin and Ross 1994; Hjelmfelt and Ross 1995; Hjelmfelt et al. 1993), where changes in concentrations are abrupt and switch from low to high concentration without an effective stable intermediate at steady-state.

A literature and/or curated database search is critical for developing an accurate logic network, but will never be sufficient to identify the underlying circuitry of an aberrant cell given the high number of genetic defects that may be present in such cells. Thus, we wondered if it was possible to infer the actual underlying network in a population of cells from experimental data routinely collected in the laboratory. To this end, we developed an approach to infer the signaling network circuitry inside a cell from experimentally measured western blot data using asynchronous Boolean logic simulation methods and a heuristic search with genetic algorithms. The objective of the approach is to identify the most likely underlying structure of a signaling network by minimizing the difference between simulated model output and experimental data collected from a series of molecular perturbation experiments. We applied the method to experimental data collected from a cell line and validated the general approach with a series of *in silico* tests, where the underlying network was known a priori.

The ability to accurately infer an intracellular network from data remains a significant and difficult problem in molecular systems biology as well as a major obstacle to signaling based combination therapies. A variety of reverse engineering approaches have been developed (Crampin et al. 2004; Kholodenko et al. 2002; Stelniec-Klotz et al. 2012). When reverse engineering a gene, signaling, or metabolic network via Boolean approaches, it is necessary to identify a logical interaction between each molecule in the network, such that the observed experimental discretized data can be explained by the model. While algorithms exist to infer Boolean networks from experimental data, including REVEAL (Liang et al. 1998), the Akutsu algorithm (Akutsu et al. 1999), and ReBMM (Saeed et al. 2012), each differs in the approach used to identify a network. REVEAL uses an information criterion to reduce the network search space consistent with the observable data. The Akutsu algorithm uses an enumerating approach to find the networks consistent with observable data. ReBMM uses a sparse search algorithm based on a Bernoulli mixture model, which is used to determine the set of Boolean networks consistent with the observable data. These and similar algorithms were largely designed to infer gene regulatory logic networks from gene expression data and assume that Boolean networks are subject to deterministic synchronous updating.

In this paper, our objective is different: we are interested in the accurate inference of the underlying structure of a signaling network from experimental data of protein activation. We also rely on a stochastic and asynchronous logic method to simulate a population level response. We use a heuristic search with genetic algorithms to identify candidate logic networks consistent with observed data collected from a set of perturbation experiments. Our approach is applied to MCF10A cells and its validity is explored with a series of *in silico* network tests.

## Materials and Methods

Here we provide a detailed description of the methods used as part of our network inference approach based on both *in vitro* and *in silico* perturbation experiments.

### Western blot analysis

MCF10A cells were serum starved overnight in a 1:1 mix of Ham’s F12 and Dulbecco’s modified Eagle medium (DMEM) with 1x Pen-Strep-Glutamine, 2.5 µg/mL fungizone, 5 µg/mL gentamycin, 10µg/mL insulin, 0.5µg/mL hydrocortisone, 0.02 µg/mL EGF from murine submaxillary gland (Sigma), and 0.1µg/mL cholera toxin. In the morning, the media was changed to a 1:1 mix of Ham’s F12 and DMEM containing no serum and no supplements, and cells were incubated for 60 min with or without 50 µM of LY294002 (Cell Signaling), a PI3K inhibitor, or 1 µM of PD0352901 (Cell Signaling), a MEK inhibitor, followed by treatment with or without 0.01 µg/mL EGF human recombinant protein (Millipore Cat. 324831) for 10 minutes. After 70 minutes, cells were harvested for protein extraction. Anti-bodies against phospho-AKT (S473 and T308), AKT, phospho-PTEN (S380), PTEN, phospho-ERK (T202/T204), ERK, phospho-Raptor (S792), Raptor, phospho-mTOR (S2448), mTOR, phospho-TSC2 (T1462) and TSC2 were obtained from Cell Signaling.

### Logic Network Simulations

All logic network simulations used an asynchronous random order update scheme with *n* replicate networks. Typically, *n*=100, unless otherwise stated. Booleannet (Albert et al. 2008) was used for all asynchronous simulations. Because a random-order asynchronous update samples many different timescales (in contrast to the uniform timescales and deterministic outcomes of synchronous update methods), it is appropriate for simulating the heterogeneity in a population of cells (Chaves et al. 2005; Garg et al. 2008; Thomas 1973; Wynn et al. 2012).

Simulations ran for a pre-defined number of time steps (long enough for a logical steady state, also known as an attractor (Thomas and D’Ari 1990), to be reached). The steady state value of all nodes can be determined explicitly in a logic model if the initial values of all nodes in the network are known *a priori*. Because it will rarely be practical to measure all molecules in realistic biological networks, our simulation approach requires only knowledge of the initial activation state of nodes explicitly perturbed in the experiment simulated (e.g., constitutively ***ON*** or ***OFF*** proteins). All nodes not given an initial value at the start of a simulation are assigned random initial values in each of the *n* replicate networks. The output of an asynchronous simulation (with *n* replicate networks) is a value between 0.0 and 1.0 for each node, indicating the probability a node is activated (**ON**) in the steady state. These probabilities are calculated directly from the average state of each node in *n* networks (Albert et al. 2008). As a direct consequence, steady state node probabilities greater than 0.0 but less than 1.0 indicate that, for the conditions tested, (a) the node is oscillating as part of a limit cycle attractor, (b) the node can be in one of two fixed point attractors, or (c) the node can be in both a limit cycle and a fixed point attractor.

### Data Discretization

For the Western blots, we chose to use data from a normal cell line (MCF10A) that had unambiguous western blot signals (that is, it was clear upon visual inspection whether a node was ***ON*** or ***OFF***). Thus, it was not necessary to explicitly define a threshold when discretizing our experimental data (referred to as expected data). However, to apply a threshold parameter when discretizing western blot data where the binary state of a measured protein is not readily inferred, semi-quantitative densitometric methods are often used.

For the simulated data, we did not consider a value of 0.4 or 0.6 to be qualitatively different from 0.5 (the value expected when exactly half of the simulated networks for a given set of conditions were ***ON*** and exactly half were ***OFF*)**. Thus, this range served as the bin for the implicit threshold of 0.5. Because our simulated values were continuous probabilities that a node was ***ON*** or ***OFF***, we also decided to define a bin for 25% ***ON*** and 75% ***ON***. While in some ways these ranges are arbitrary, empirically we found that this helped our simulations converge to optimal solutions (when a “true” target network was known), suggesting that the approached helped smooth the stochastic nature of these probability values.

More details about the data discretization can be found in the Supplementary Materials of Wynn et al. (2014).

### Predictive Scores

Comparison of model readout with experimental data necessitated the discretization of experimental western blot data. It is important to note that because a phospho western blot is a qualitative assay dependent on many factors (including film exposure time), attempts at precise quantitation may be misleading and may not be necessary for the modeling approach used here. Values of ***ON*** or ***OFF*** were assigned to indicate the phosphorylation state of proteins in MCF10A cells for each condition tested. In general, faint or absent blots were assigned ***OFF*** values (0.0) and prominent blots were assigned ***ON*** values (1.0).

A predictive score (p-score) was developed to quantify how well simulated model output matched experimentally collected data (referred to as expected values). The p-score is calculated as:

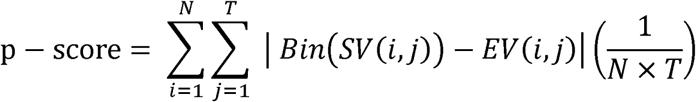
where, *i* is the node identifier, *j* is the test condition identifier, *N* is the total number of nodes evaluated in the network, *T* is the total number of test conditions, *SV*(*i, j*) is the simulated value of the *i*-th node in the *j*-th test condition, *EV*(*i, j*) is the expected value of the i-th node in the j-th test condition. The function *Bin*(*x*), partitions simulated values as follows (input → output):

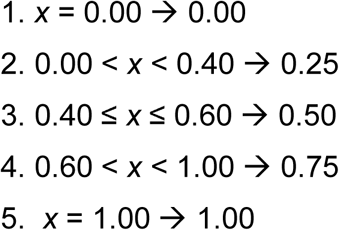
The *N* nodes evaluated in the p-score include only those nodes for which expected values were available. Consequently, *N* may be smaller than the total network node count. The scaled p-score will range from 0.0 (indicating a perfect match with experimental data) to 1.00 (indicating that the simulated output was wrong for all evaluated nodes in all test conditions).

### Genetic Algorithm and Fitness Function

An implementation of a genetic algorithm (GA) was used to identify candidate logic networks from a large search space of possible logic networks. A GA is a type of heuristic search that is often used to find optimal solutions in very large search spaces that cannot reasonably be searched exhaustively. GAs leverage operations that mimic aspects of biological evolution to identify an optimal solution, such as fitness selection, reproduction, chromosomal crossover, and random mutation (Crampin et al. 2004; Deb 1999; Mitchell 1996). While GA operations may appear simple, the search performed is considered “highly nonlinear, massively multi-faceted, stochastic, and complex” (Deb 1999). Typically, the problem to be optimized is numerically encoded via vectors of numbers referred to as chromosomes. A fundamental component of a GA is a fitness function that assigns a numerical value based on the relative fitness of each chromosome. A GA begins with an initial population of chromosomes and proceeds by allowing the population to evolve over many generations.

We chose to use a GA because the search space in which we were looking for an optimal network configuration was often far too large to search exhaustively. In our case, the GA attempts to minimize the difference between experimental input data and simulated output of candidate logic networks under the same perturbation conditions used to collect the experimental data (**Figure 1**). The output of the fitness function used is a p-score. Our GA implementation does not stop when an optimal solution is found (i.e., a network with a p-score of 0.0) because we want the population to continue to evolve to other local optima, if they exist. The general steps followed in our GA implementation are:

1. In generation 0, randomly initialize a population of *p* chromosomes as a numeric vector, where each element of the vector corresponds to a node in the network, and each numeric value corresponds to a possible logic regulation rule for the node. Each unique chromosome represents a unique numeric encoding of a candidate logic network.
2. In each generation, evaluate the chromosomes in the current population by calculating their p-scores via a fitness function. The next generation’s population is determined by cross-over via random tournament selection (Mitchell 1996) and mutation rates between 0.2 and 0.5.
3. Report all networks with an optimal p-score of 0.0.
4. Run until the number of generations set at the start of the simulation is reached.

**Figure 1.**
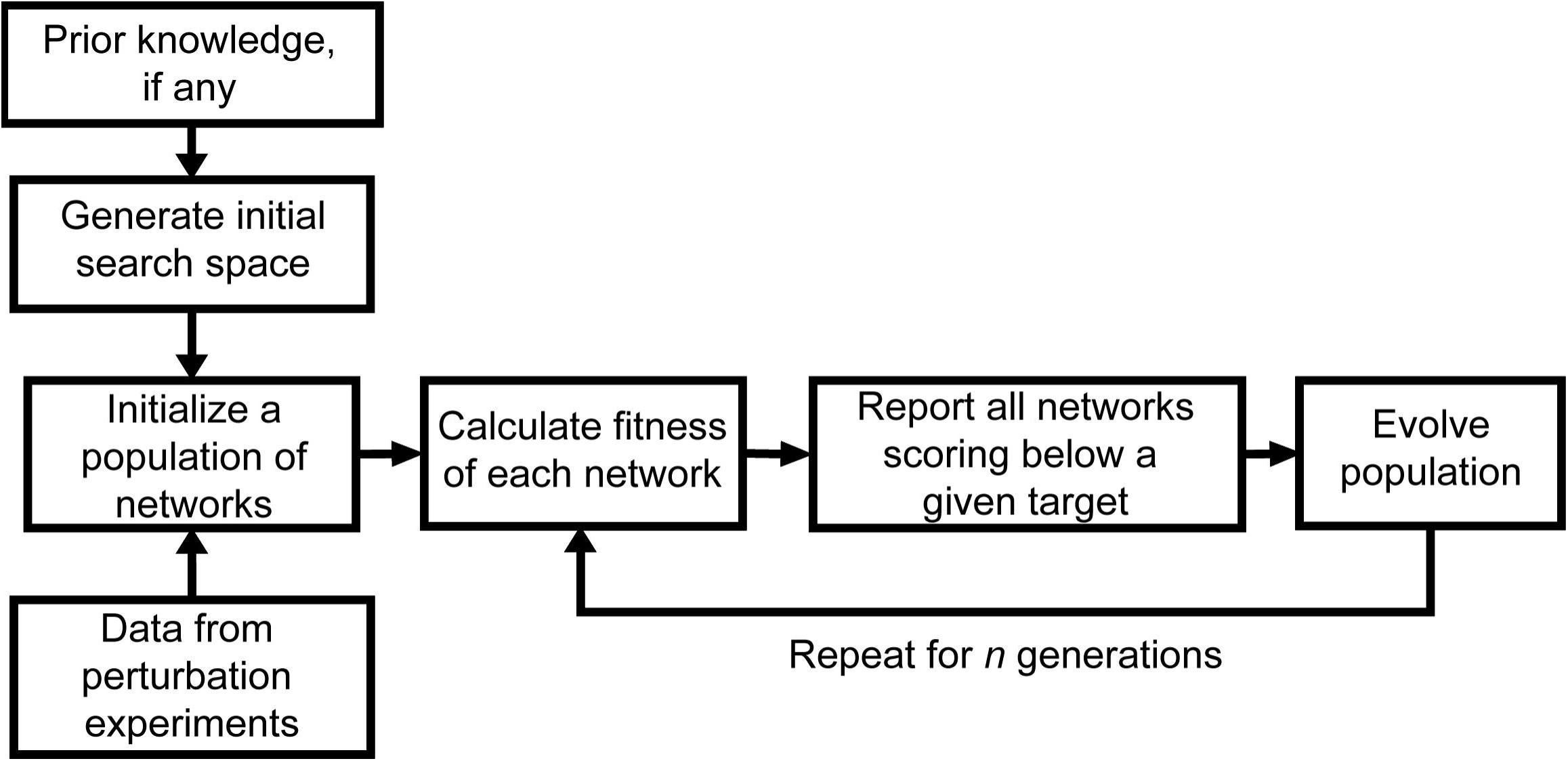
High-level flow chart of the proposed methodology with genetic algorithm. Data from perturbation experiments are determined into an ON or OFF value and served as an input for the fitting. Prior knowledge on the network based on literature search, if any, defined the input initial search space. The genetic algorithm generates and evolves populations of networks based on the calculated fitness to the input experimental data. The algorithm evolves the output networks and calculates the fitness for the user-defined *n* generations.

For very large search spaces, GA were performed three times for at least 100 generations each to ensure that adequate coverage of the large search space was achieved. If there is uncertainty about the coverage of the search space, the number of generations can be increased in our approach.

### Logic network search space generation

The space of all possible logic networks searched by the GA consists of all logic network combinations that can be made for the regulations possible for each node in a network. A script was used to enumerate all logically unique ways a single node could be regulated by all or a subset of nodes in a network (depending on whether any prior knowledge of the network is available). For *m* regulatory nodes, which represent the maximal set of nodes that can possibly regulate a target node, we assume each regulatory node appears no more than one time in a logic rule. Because order matters in a logical expression, we generate all permutations of the ways 1 to *m* nodes can by linked by logic operators as well as by parenthetic groupings. Finally, we scan all generated rules to check for logical equivalency and remove any redundant expressions (that is, any expression that generates a truth table identical to the truth table of an expression already included in the set). A unique identifier is assigned to each logically unique regulation rule possible for a target node. **Table 1** lists the number of unique logic regulations possible when a node is regulated by up to 6 other nodes in the network. **Table 2** and **Supplemental Table 1** show examples of logic regulations that make up two distinct logic network search spaces.

**Table 1.** The complexity of the network search space grows with the number of nodes in the network.

| Nodes in network ($n$) | 2 | 3 | 4 | 5 | 6 |
| --- | --- | --- | --- | --- | --- |
| maximum number of regulator nodes per node ($n-1$) | 1 | 2 | 3 | 4 | 5 |
| interaction networks possible ($3^{n(n-1)}$) | $3^2 = 9$ | $3^6 = 729$ | $3^{12} = 531,441$ | $3^{20} = 3.5 \times 10^9$ | $3^{30} = 2.1 \times 10^{14}$ |
| maximum unique logic regulation rule count for a single node ($r$) | 3 | 13 | 95 | 1145 | 20,825 |
| maximum logic network count ($r^n$) | $3^2 = 9$ | $13^3 = 2,197$ | $95^4 = 1.8 \times 10^{11}$ | $1,145^5 = 2.0 \times 10^{15}$ | $20,825^6 = 8.2 \times 10^{25}$ |

**Table 2.** All possible logic rules by node in the informed search space of the hypothesized PI3K and MAPK signaling network. The 43 rules combine to form an informed search space of 41,067 possible networks, which is the product of the regulation rule count of each node. All rule identifiers equal to 0 correspond to the case where the node is an input node (and no other node in the network regulates its activation state).

| rule count | node rule identifier | node | regulation rule |
| --- | --- | --- | --- |
| 1 | 0 | AKT | $AKT^{t+1} = AKT^t$ |
| 2 | 1 | AKT | $AKT^{t+1} = \text{not } AKTS473^t$ |
| 3 | 2 | AKT | $AKT^{t+1} = AKTS473^t$ |
| 4 | 3 | AKT | $AKT^{t+1} = \text{not } AKTT308^t$ |
| 5 | 4 | AKT | $AKT^{t+1} = AKTT308^t$ |
| 6 | 5 | AKT | $AKT^{t+1} = \text{not } AKTS473^t \text{ and not } AKTT308^t$ |
| 7 | 6 | AKT | $AKT^{t+1} = \text{not } AKTS473^t \text{ or not } AKTT308^t$ |
| 8 | 7 | AKT | $AKT^{t+1} = \text{not } AKTS473^t \text{ and } AKTT308^t$ |
| 9 | 8 | AKT | $AKT^{t+1} = \text{not } AKTS473^t \text{ or } AKTT308^t$ |
| 10 | 9 | AKT | $AKT^{t+1} = AKTS473^t \text{ and not } AKTT308^t$ |
| 11 | 10 | AKT | $AKT^{t+1} = AKTS473^t \text{ or not } AKTT308^t$ |
| 12 | 11 | AKT | $AKT^{t+1} = AKTS473^t \text{ and } AKTT308^t$ |
| 13 | 12 | AKT | $AKT^{t+1} = AKTS473^t \text{ or } AKTT308^t$ |
| 14 | 0 | AKTS473 | $AKTS473^{t+1} = AKTS473^t$ |
| 15 | 1 | AKTS473 | $AKTS473^{t+1} = \text{not } PI3K^t$ |
| 16 | 2 | AKTS473 | $AKTS473^{t+1} = PI3K^t$ |
| 17 | 0 | AKTT308 | $AKTT308^{t+1} = AKTT308^t$ |
| 18 | 1 | AKTT308 | $AKTT308^{t+1} = \text{not } PIP3^t$ |
| 19 | 2 | AKTT308 | $AKTT308^{t+1} = PIP3^t$ |
| 20 | 0 | EGFR | $EGFR^{t+1} = EGFR^t$ |
| 21 | 0 | ERK | $ERK^{t+1} = ERK^t$ |
| 22 | 1 | ERK | $ERK^{t+1} = \text{not } MEK^t$ |
| 23 | 2 | ERK | $ERK^{t+1} = MEK^t$ |
| 24 | 0 | MEK | $MEK^{t+1} = MEK^t$ |
| 25 | 1 | MEK | $MEK^{t+1} = \text{not } EGFR^t$ |
| 26 | 2 | MEK | $MEK^{t+1} = EGFR^t$ |
| 27 | 0 | PI3K | $PI3K^{t+1} = PI3K^t$ |
| 28 | 1 | PI3K | $PI3K^{t+1} = \text{not } EGFR^t$ |
| 29 | 2 | PI3K | $PI3K^{t+1} = EGFR^t$ |
| 30 | 0 | PIP3 | $PIP3^{t+1} = PIP3^t$ |
| 31 | 1 | PIP3 | $PIP3^{t+1} = \text{not } PI3K^t$ |
| 32 | 2 | PIP3 | $PIP3^{t+1} = PI3K^t$ |
| 33 | 3 | PIP3 | $PIP3^{t+1} = \text{not } PTEN^t$ |
| 34 | 4 | PIP3 | $PIP3^{t+1} = PTEN^t$ |
| 35 | 5 | PIP3 | $PIP3^{t+1} = \text{not } PI3K^t \text{ and not } PTEN^t$ |
| 36 | 6 | PIP3 | $PIP3^{t+1} = \text{not } PI3K^t \text{ or not } PTEN^t$ |
| 37 | 7 | PIP3 | $PIP3^{t+1} = \text{not } PI3K^t \text{ and } PTEN^t$ |
| 38 | 8 | PIP3 | $PIP3^{t+1} = \text{not } PI3K^t \text{ or } PTEN^t$ |
| 39 | 9 | PIP3 | $PIP3^{t+1} = PI3K^t \text{ and not } PTEN^t$ |
| 40 | 10 | PIP3 | $PIP3^{t+1} = PI3K^t \text{ or not } PTEN^t$ |
| 41 | 11 | PIP3 | $PIP3^{t+1} = PI3K^t \text{ and } PTEN^t$ |
| 42 | 12 | PIP3 | $PIP3^{t+1} = PI3K^t \text{ or } PTEN^t$ |
| 43 | 0 | PTEN | $PTEN^{t+1} = PTEN^t$ |

### Computing environment

A custom python library called Netinf was written to support the network inference search and implementation of the GA. The library, which makes calls to Booleannet (Albert et al. 2008) for asynchronous network simulations, supports distributed computing using MPI for Python. All distributed simulations were run on the University of Michigan’s high performance computing cluster using Intel Nehalem/i7 Cores.

## Results

The objective of our approach presented in this study is to accurately infer the underlying structure of a signaling network from experimental data. We first consider a hypothesized network of EGF based signaling in MCF10A cells. We then evaluate our method by testing its ability to identify a series of increasingly complex test networks.

### EGFR signaling in MCF10A cells

A detailed literature search was performed to identify the important regulatory components of EGF induced signaling in the PI3K and MAPK pathways (**Table 3**). From this information, a canonical normal interaction network was constructed (**Figure 2A**). Interaction networks provide directional information about a set of regulatory events, but do not provide details about the hierarchy of regulations when more than one node regulates another node (Vidal et al. 2011; Wynn et al. 2012). For example, the interaction network indicates that **PIP3** is activated by **PI3K** and inhibited by **PTEN**, but does not indicate which effect is dominant when both molecules are present. The literature supporting each regulation in the interaction network was, therefore, carefully considered before deriving a set of hypothesized logic rules (**Table 4**), which make up the hypothesized logic network of normal EGF signaling in MCF10A cells (**Figure 2B**). In general, an ***AND NOT*** regulation means the inhibitor will dominate when both an activator and inhibitor are present, while an ***OR NOT*** regulation means that the activator will dominate when both an activator and inhibitor are present [see (Wynn et al. 2012) for a more detailed discussion].

**Table 3.**
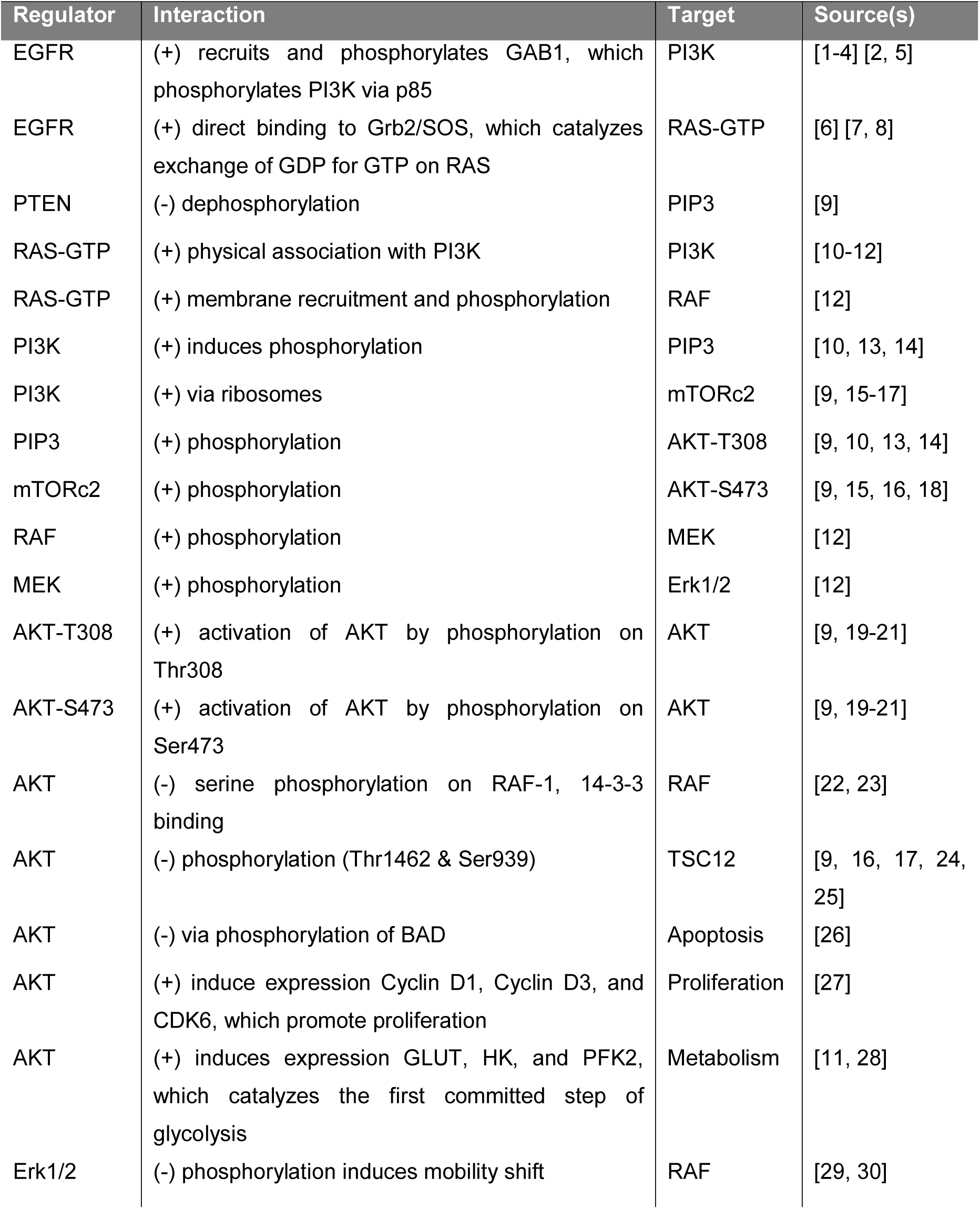
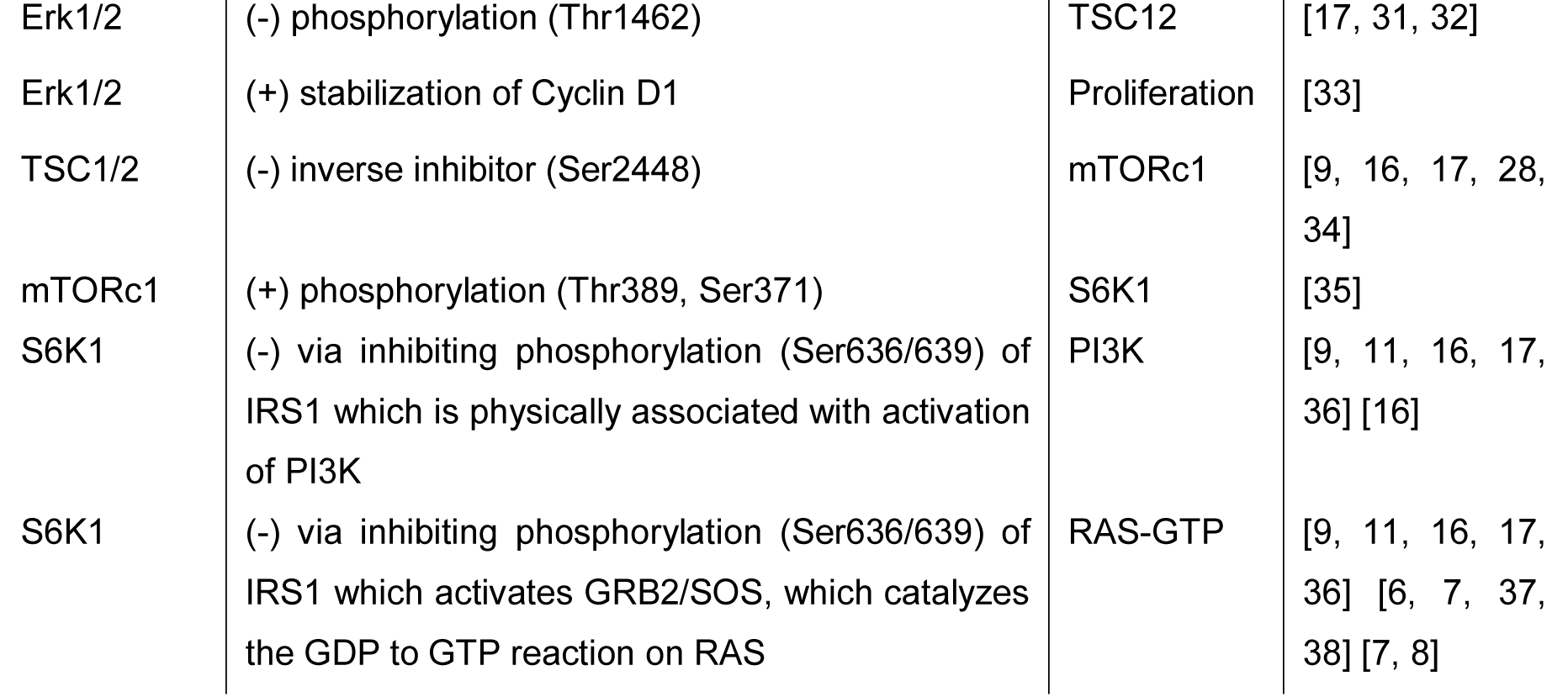
Table of key interactions in the PI3K and MAPK signaling pathways. The (+) and (-) symbols indicate activation and inhibition, respectively, of a target molecule by a regulator molecule. Source references are available in the Appendix.

**Figure 2.**
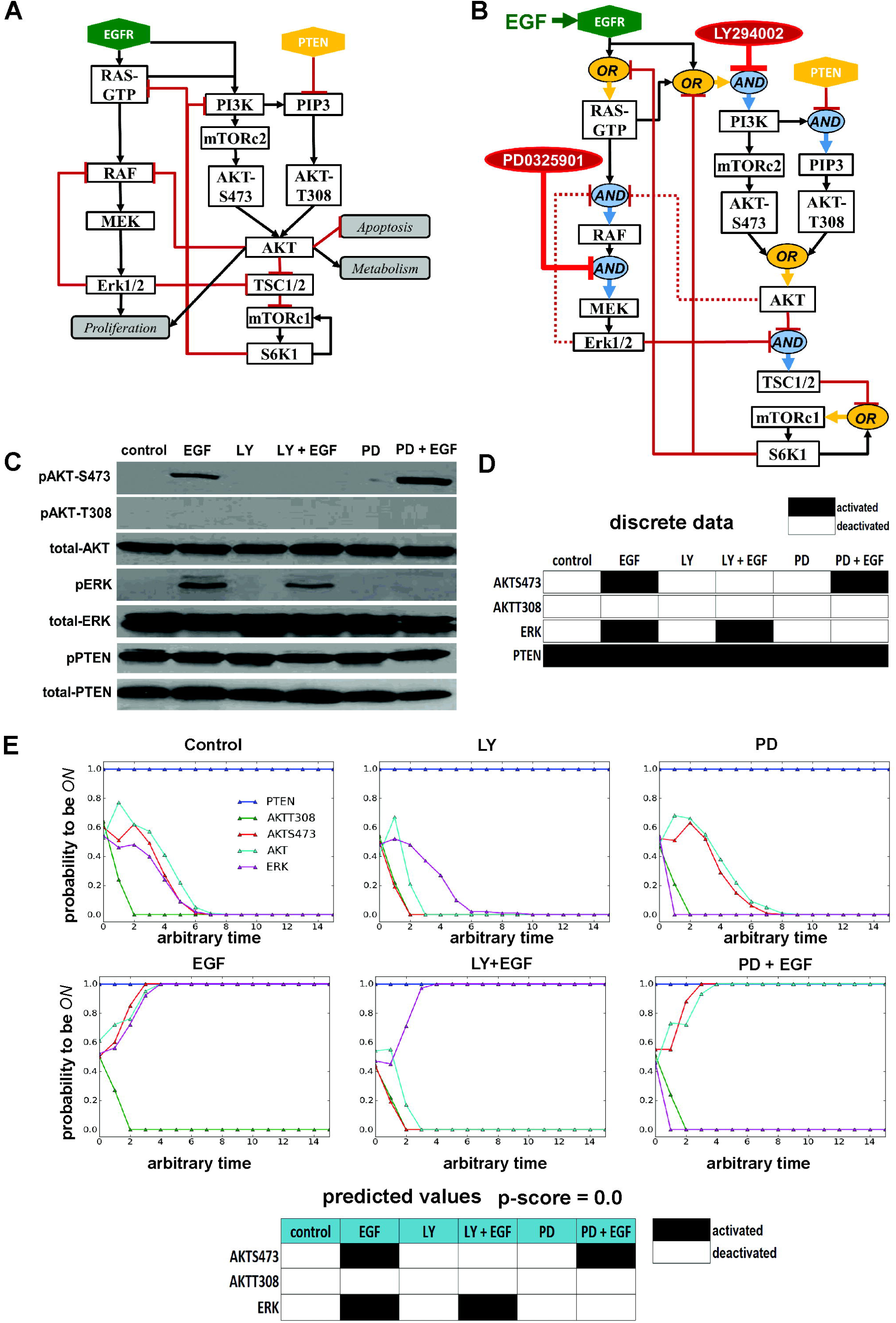
Hypothesized network of epidermal growth factor receptor signaling in MCF10A cells. The hypothesized **(A)** interaction and **(B)** logic networks of normal-like EGF-driven signaling of the PI3K and MAPK pathways in MCF10A cells are presented. Logic network simulations included 15 nodes: **EGFR, PTEN, RAS-GTP, RAF, TSC12, PIP3, PI3K, S6K1, mTORc1, mTORc2, MEK, AKT, AKTT308, AKTS473, and ERK.** (**C**) Western blots of total and phosphorylated AKT, ERK, and PTEN in MCF10A cells in six experimental perturbation conditions are presented along with (**D**) the discretized form of these data as logical expected values. (**E**) The asynchronously simulated output of the hypothesized logic network for each perturbation condition as well as predicted steady states as logical values are also presented. The hypothesized logic network produced a p-score of 0.0.

**Table 4.** Logic function justifications for the hypothesized normal logic network of the PI3K and MAPK pathways in MCF10A cells. Source references are available in the Appendix.

| Logic Rule | Rule Justification | Source(s) |
| --- | --- | --- |
| $PTEN^{t+1} = PTEN^t$ | PTEN is an input to the model. Phosphorylated PTEN is always present under the conditions tested in MCF10A cells | [39] |
| $EGFR^{t+1} = EGFR^t$ | EGF is an input to the model, which stimulated its receptor, EGFR. The receptor is assumed to be activated only when EGF is added to MCF10A cells. | [40] |
| $AKT^{t+1} = AKT-T308^t \text{ OR } AKT-S473^t$ | While AKT requires phosphorylation at both AKT-T308 and AKT-S473 for full activation, we assume that partial activation is possible when only one is present. In support of this are data indicating that cells expressing PTEN were still able to activate AKT. | [20] |
| $AKT-S473^{t+1} = mTORC2^t$ | There is evidence that mTORC2 is essential for full AKT activation (by directly phosphorylating AKT, likely due to PDK2 activity). We assume PDK2 is present in the system but do not include it in the model for simplicity. | [9, 19] |
| $AKT-T308^{t+1} = PIP3^t$ | PIP3 recruits AKT to the cell membrane where AKT is activated by PDK1. We assume PDK1 is present in the system but do not include it in the model for simplicity. | [9, 20, 21] |
| $RAF^{t+1} = RAS-GTP^t \text{ AND NOT } AKT^t \text{ AND NOT } Erk1/2RK^t$ | RAS-GTP directly activates RAF, and RAF must be phosphorylated and dephosphorylated on specific sites to remain active. Regardless of the activation state of RAF, AKT appears able to phosphorylate RAF, allowing the small molecule 14-3-3 to bind and reduce activation levels. Therefore, the rule is written as RAS-GTP 'and not AKT'. In addition, Erk1/2 is able to phosphorylate RAF on specific sites that induce a mobility shift in RAF, therefore reducing activation levels. The interaction between RAF and Erk1/2 appears independent of AKT, therefore the rule is written with 'and not Erk1/2'. | [22, 23]<br>[12] [29, 30] |
| $Erk1/2^{t+1} = MEK^t$ | ERK1/2 is directly activated by MEK in the MAPK pathway | [41] |
| $MEK^{t+1} = RAF^t$ | RAF is upstream activator of MEK. The MEK inhibitor, PD0325901, is assumed to override activation of MEK by RAF phosphorylation in all conditions tested. The inhibitor is not modeled as a separate node in the network. | [41, 42]<br>[43] |
| $mTORC1^{t+1} = S6K1^t \text{ OR NOT } TSC12^t$ | S6K1 has a positive feedback, activating mTORC1 via h6a and h6c short isoforms. TSC1 and TSC2 form a heterodimer that inhibits the activity of Rheb. Rheb is a small GTPase required for activation of mTORC1. Thus, in the model we assume that mTORC1 is activated by S6K1 positive feedback or inactivated by TSC1/2 activity. | [44] [9] |
| $mTORc2^{t+1} = PI3K^t$ | Evidence strongly suggests that PI3K is both necessary and sufficient for mTORc2 activation to occur via mTORc2 ribosomal binding. We assume that the ribosome concentration is above the regulatory threshold in the cells and therefore consider mTORc2 activation regulated by only PI3K. | [45, 46] |
| $PI3K^{t+1} = EGFR^t$<br>**OR** RAS-GTP<sup>t</sup> **OR**<br>**NOT** S6K1<sup>t</sup> | This is a particularly complicated regulation. We simplify the network by using EGFR as a direct activator of PI3K instead of detailing the chain of activation from EGFR to GAB1, which normally activates PI3K by recruiting the p85 subunit. Erk1/2 negatively regulates PI3K by phosphorylating GAB1 and rendering it unable to recruit p85, but we assume this negative feedback is not activated within an hour and therefore do not include the feedback in the regulation of PI3K. In addition, S6K1 inhibits IRS1, which activates PI3K. Because insulin, an activator of IRS1, is not added in our experiments, we assume that IRS1 is only regulated by S6K1. Therefore, we simplify the interaction by writing S6K1 as an inhibitor of PI3K, independent of EGFR. Finally, RAS-GTP can also activate PI3K and its activation is not affected by EGFR or S6K1 levels. Finally, we assume the PI3K inhibitor, LY294002, overrides all activators of PI3K and results in complete inhibition of PI3K. The inhibitor is not modeled as a separate node in the network. | [1, 2, 5, 10, 16, 36, 47-52] [53] |
| $PIP3^{t+1} = PI3K^t$<br>**AND NOT** PTEN<sup>t</sup> | PI3K activates PIP3 at the cell membrane and PTEN dephosphorylates PIP3. We assume the inhibitor is dominant. | [9] |
| $RAS-GTP^{t+1} = EGFR^t$ **OR** **NOT**<br>S6K1<sup>t</sup> | EGFR stimulates RAS-GTP via a GRB2/SOS complex, assumed to be present in all cells. Furthermore, S6K1 inhibits IRS1, which activates RAS-GTP. For simplicity, we leave IRS1 out of our network, as it is assumed to be only regulated by S6K1. | [1, 12] [36] |
| $S6K1^{t+1} = mTORc1^t$ | Phosphorylation of S6K1 is induced by mTORc1. | [54] |
| $TSC1/2^{t+1} = NOT$<br>AKT<sup>t</sup> **AND** **NOT**<br>Erk1/2<sup>t</sup> | AKT inhibits TSC1/2 and Erk1/2 inhibits TSC1/2 through the activation of RSK1. Phosphorylation (Ser1798) of RSK1 inhibits TSC1/2 and results in increased mTORc1 expression. We assume that TSC1/2 is only **ON** when its activator is present and both its inhibitors are absent. While there are a number of negative regulators of the TSC1/2 dimer, our model only considers Erk1/2 inhibition and AKT inhibition of TSC1/2. | [32] |

The correctness of the hypothesized logic network model (**Figure 2B**) for normal-like cells was tested by comparing *in silico* model predictions to experimentally measured readouts of phosphorylated AKT (at S473 and T308) and ERK (at T202/T204) in the normal-like mammary epithelial MCF10A cell line. Six experimental conditions were performed *in vitro* (**Figure 2C**) and *in silico* (**Figure 2D**): (1) control, (2) EGF stimulation, (3) LY294002 (LY) alone, (4) LY followed by EGF stimulation, (5) PD0325901 (PD) alone, and (6) PD followed by EGF stimulation. For *in vitro* immunoblot experiments, cells were serum starved overnight and then treated with or without PD, a direct inhibitor of MEK (Leyton et al. 2008), or LY, a direct inhibitor of PI3K (Qiang et al. 2004) for 1 h, followed by 10 min with or without EGF stimulation. For *in silico* experiments, the control condition was simulated by fixing the **EGFR** node to ***OFF*** for the duration of the simulation, while the presence of EGF in test conditions was simulated by fixing the **EGFR** node to ***ON*** for the duration of the simulation. The presence of LY or PD in test conditions was simulated by fixing **PI3K** or **MEK**, respectively, to ***OFF*** for the duration of the simulation. In all six experimental conditions tested, PTEN (which is known to be wild-type in MCF10A cells) was activated (**Figure 2C**). As a consequence, the **PTEN** node was fixed to ***ON*** in all simulations of MCF10A cells.

The hypothesized logic network (**Figure 2B**) correctly predicted the phosphorylation of **AKTS473**, **AKTT308**, and **ERK** in all six experimental conditions (**Figure 2C**), producing a p-score of 0.0, which indicates an exact qualitative match between experiment and simulation (see **Methods**). The hypothesized logic model does not include inhibitory regulation of **RAF** by **AKT** or **ERK** (Brummer et al. 2003; Ritt et al. 2010; Rommel et al. 1999; Zimmermann and Moelling 1999) (as indicated by dashed red lines in **Figure 2B**). We hypothesized that the negative regulation of **RAF** by **AKT** and **ERK**, which would be the only sources of **ERK** oscillations in our model, are “late effects” (as per the term used by Samaga et al. (2009) that will take place much later than our experimental timescale of 70 minutes. A longer experiment with multiple time points would be required to effectively measure possible oscillatory changes in ERK activation. Some experiments that report rapid growth factor induced oscillations in phosphorylated ERK, report changes in the relative ratio of phospho-ERK to total-ERK (Nakayama et al. 2008; Shankaran et al. 2009). While the ratio may be changing, a robust phospho ERK signal is often still observed. When an alternate version of the network that included negative regulation of **RAF** by **AKT** and **ERK** was tested, it produced a p-score of 0.08 (see **Supplemental Figure 1**).

These results suggest that the hypothesized model (**Figure 2B**) may represent the underlying circuitry driving EGF based signaling in the PI3K and MAPK pathways in MCF10A cells, but do not definitively establish this to be the case. In an attempt to confirm or refute the proposed circuitry, we wondered how many other logical configurations of the nodes in the hypothesized logic network model could generate a p-score of 0.0. While an exhaustive search of all logic network configurations was not practical for a network of this size (**Table 1**), an optimized search via a GA (see **Methods**) found hundreds of alternate logic network configurations that can also generate a p-score of 0.0 based on the expected values of **AKTS473**, **AKTT308**, and **ERK** (**Figure 3C**). A subset of these networks can be discarded because the regulations involved do not represent what is expected in a normal model of the pathways considered. For example, all networks with a p-score of 0.0 that include PTEN activating (rather than inhibiting) **PIP3**’s role in the phosphorylation of **AKTT308** could be rejected. It may be imprudent to make this assumption, however, if we were attempting to uncover the signaling network inside a cancer or aberrant cell with unknown signaling adaptations.

**Figure 3:**
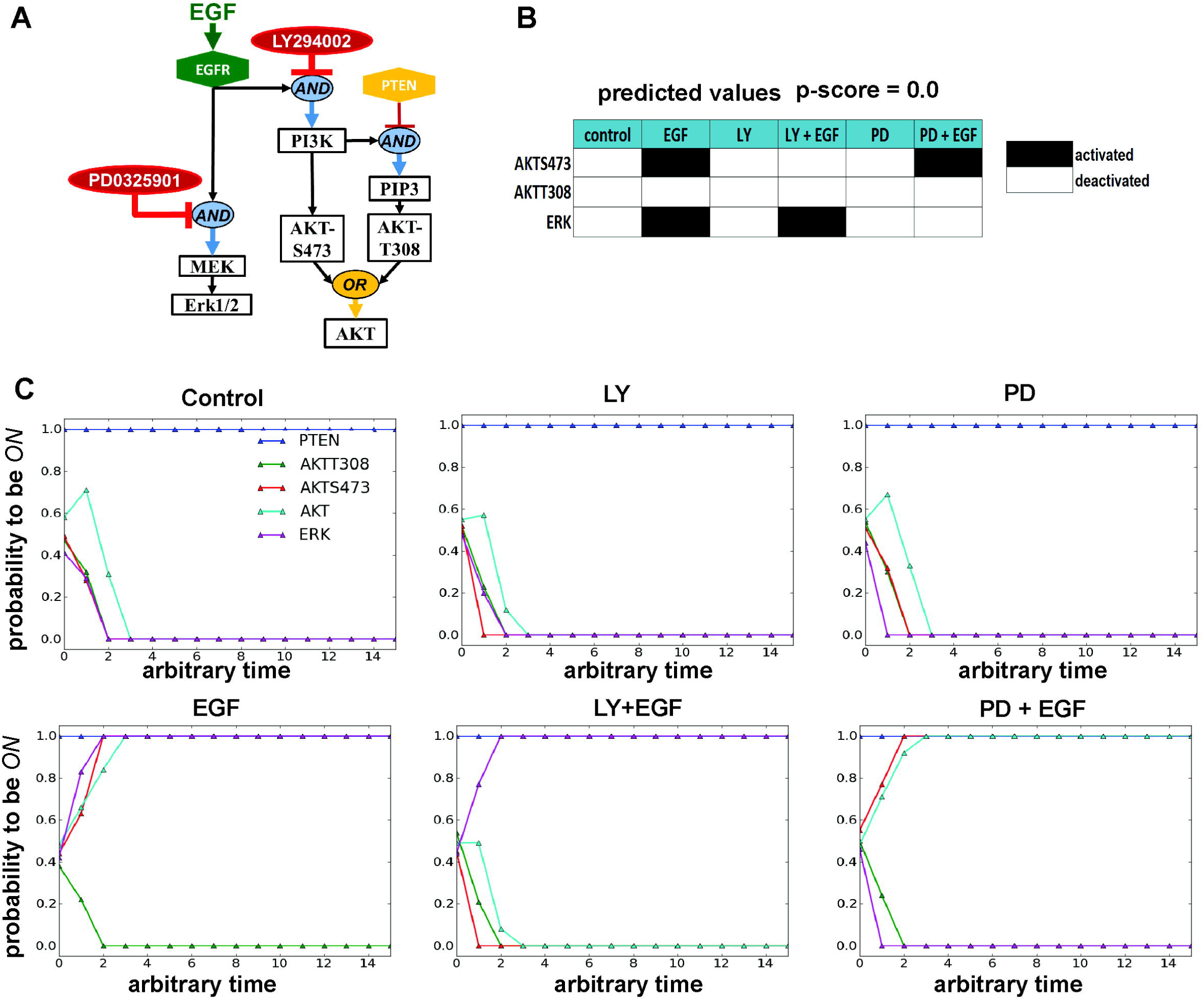
Simplified form of the hypothesized network of epidermal growth factor receptor signaling in MCF10A cells. The hypothesized logic network was simplified to 9 nodes (**EGFR, PTEN, PIP3, PI3K, MEK, AKT, AKTT308, AKTS473, and ERK**). The steady-state output of this network is qualitatively identical to the larger hypothesized network (**Figure 2B**). The predicted steady state values (p-score = 0.0) and asynchronous simulation results under the six perturbation conditions are presented.

To test if including experimental measurements for all or most nodes in the hypothesized network will narrow the list of candidate networks, we constructed a smaller 9-node hypothesized logic network (**Figure 3A**) where **PIP3** was the only node not directly measured or perturbed experimentally. While this version of the logic network also generated a p-score of 0.0 (**Figure 3B**), it is still not clear that the reduced network is an accurate representation of the underlying EGF based signaling circuitry of MCF10A cells. If we know which nodes *can* regulate other nodes in the network with a high degree of accuracy, but do not know precisely how these regulations occur, it is possible to reduce the complexity of the search space in which to look for other networks that may have a p-score of 0.0. First, **EGFR** and **PTEN** can be assumed to be input nodes (which means no other node in the network directly regulates them). This assumption is justified for PTEN because the phosphorylation of PTEN is unchanged in the 6 experimental conditions tested (**Figure 2C**). From the interaction network (**Figure 2A**), it can be assumed that **MEK** is regulated by **EGFR**, **ERK** is regulated by **MEK**, **PI3K** is regulated by **EGFR**, **PIP3** is regulated by **PI3K** and/or **PTEN**, and **AKT** is regulated by some combination of the phosphorylation of its T308 and S473 residues (which in turn can be assumed regulated by **PIP3** and **PI3K**, respectively).

If we assume only that the above regulations are possible, but assume nothing about the exact nature of the regulations, then a total of 36 logic regulation rules are possible. If the possibility that nodes other than **EGFR** and **PTEN** could be input nodes is included (in the case of **AKTS473** and **AKTT308** this would allow for the possibility of auto-phosphorylation), then 43 distinct logic regulation rules are possible (**Table 2**), which combine to form a search space of 41,067 distinct logic networks. p-scores were calculated after simulating all 41,067 logic networks, and 78 networks were found with a p-score of 0.0, leaving additional room for doubt about whether our hypothesized network (**Figure 3A**) can be considered correct.

Returning to the original hypothesized interaction network (**Figure 2A**), there are 3^210^ (3^n^(^n-1^), n = 15) possible interaction networks that can be constructed from the 15 nodes in this network and many more logic networks. If we make assumptions similar to those we made for the reduced network, there are a total of 275 possible logic rules that can regulate the 15 nodes, producing a very large search space of **2.4 x 10^12^** logic networks (**Supplemental Table 1**). Obtaining additional expected values by performing additional perturbation experiments will likely improve the probability of finding the network that best reflects the network circuitry in the cell. Because laboratory experiments are resource intensive, it would be advantageous to be able to predict the next set of experiments that will best inform the network search. To this end, we designed a series of *in silico* experiments to test whether it is possible to infer the underlying signaling network from experimental data using a series of increasingly complex *in silico networks*. We also tested whether it is possible to reasonably predict the next best set of experiments to perform in the lab to facilitate discovery of the network that best represents the underlying signaling circuitry of a given cell type.

### Validation of methodology in silico

We evaluated our network inference methodology with a series of test networks. We assume that between any two nodes in a logical circuit there can be, at most, two directional edges (depicted as a black dashed line in **Figure 4A**), where each directional edge can be in one of three regulatory states: (0) absent with no regulation, (1) inhibiting the target node, or (2) activating the target node. In a two-node network the number of interaction networks and logic networks are equivalent (because there are no ***AND*** or ***OR*** operators possible). For *n* > 2, however, the number of logic networks grows faster than the number of possible interaction networks (**Table 1**).

**Figure 4.**
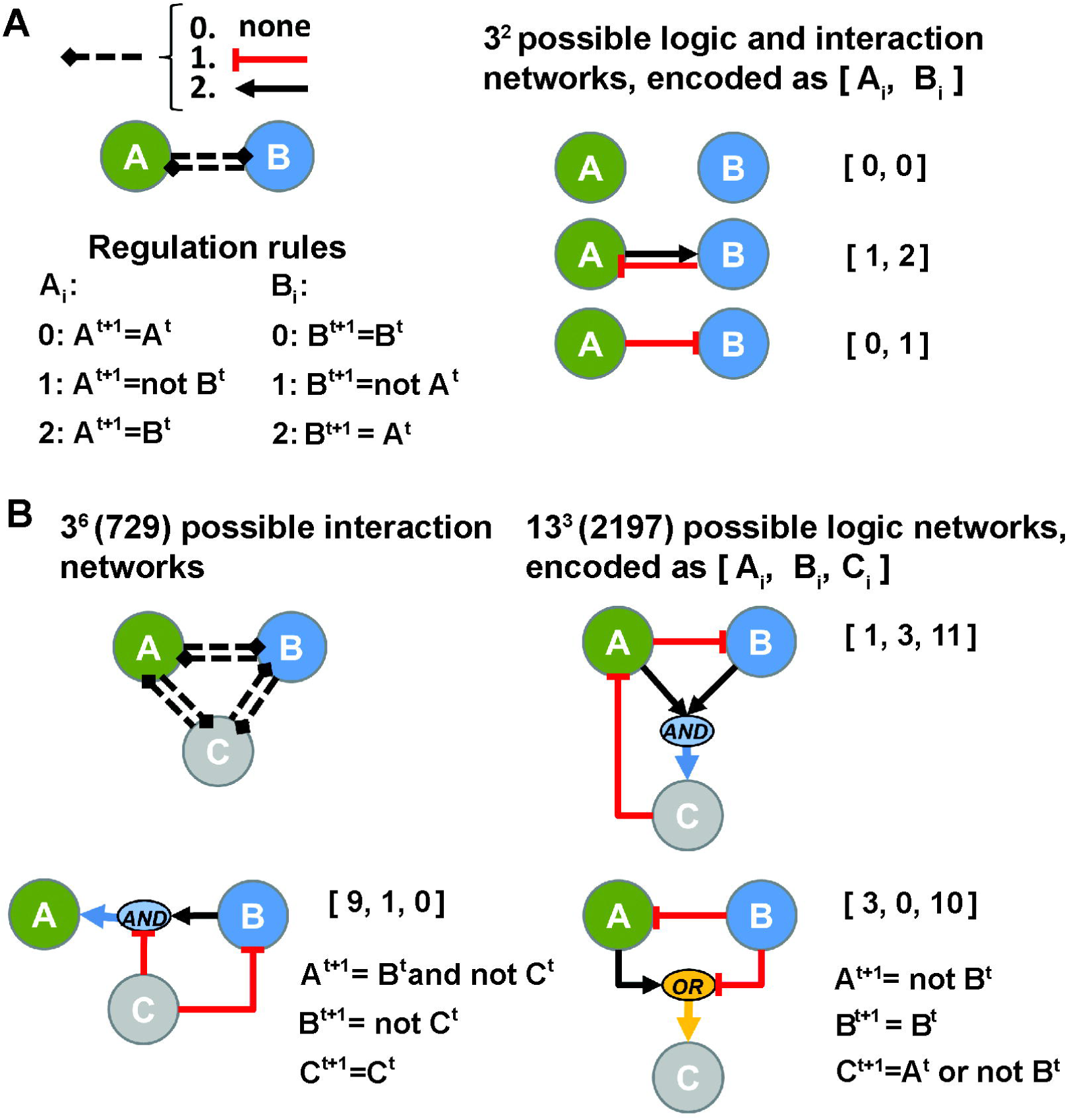
Simple logic networks. **(A)** Each black dashed edge between two nodes is a directional edge representing one of three regulation states: (0) no regulation, (1) inhibition, or (2) activation. For a two node network consisting of **A** and **B**, the logic rules corresponding to each node are listed. A total of 9 logic networks can be constructed from these rules. Each logic network can be uniquely encoded as **[A_i_, B_i_]**, where **A_i_** and **B_i_** correspond to one of the three regulation rules possible for each node. For example, **[0, 0]** represents the case where no regulation exists between either node (i.e., they are both input nodes), **[1, 2]** represents the case where **A** is inhibited by **B** and **B** is activated by **A**, and **[0, 1]** represents the case where **A** is an input node and **B** is inhibited by **A**. **(B)** Similarly, for a three-node network consisting of **A, B**, and **C**, each logic network can be uniquely encoded as **[A_i_, B_i_, C_i_,].** The logic rules corresponding to each node are listed in **Supplemental Table 2.** A total of 729 interaction networks and 2,197 logic networks are possible for these three nodes

The objective of the *in silico* tests performed was to recover the correct network from a large search space of networks (that is, to find a single network with a p-score of 0.0) using only the expected values generated by the correct network from a set of *in silico* perturbation experiments. The general approach followed was:

1. For all nodes in a test network known a priori, create a network search space by generating the set of all possible logic rules that may regulate each node (this set may be reduced by prior knowledge).
2. Identify a set of network perturbation experiments to perform *in silico*.
3. Run simulations of the test network under each perturbation condition to generate expected values.
4. Search for logic network configurations that can produce the same expected values as the target test network did in step 3, under identical perturbation conditions (i.e., p-score = 0.0). For large search spaces, the search was optimized using a GA, where the fitness function generates a p-score (see **Methods**) for each candidate network in the population of each generation.
5. If multiple networks with a p-score of 0.0 are found, attempt to reduce this number by predicting additional perturbation experiments that can differentiate between these networks. Repeat step 3.

### Two-node test network

In a two-node network consisting of **A** and **B**, a total of 9 possible logic networks are possible. These networks can be uniquely identified in vector form as **[A_i_, B_i_]**, where **A_i_** and **B_i_** correspond to one of the three logic rules possible for each node (**Figure 4A**). In this very simple example, the number of interaction and logic networks are equivalent (**Table 1**). For each of the 9 logic networks that can be formed from **A** and **B**, expected values were generated *in silico* with the following perturbations: (1) **A *ON***, (2) **A *OFF***, (3) **B *ON***, and (4) **B *OFF*** (**Supplemental Figure 2**). The *in silico* data were used as expected values analogous to how the *in vitro* western blot data was used as expected values for the MCF10A cells (Figure 2). For each of the 9 sets of expected values, we ran a search that attempted to recover each of the 9 original networks. Because the search space is small, it was straightforward to perform an exhaustive search. As an example, when network **[1, 2]** was the target search network (**Figure 5**), a p-score was generated for all 9 candidate networks. In all 9 searches, the correct network was the only network with a p-score of 0.0 and, thus, the correct network was always recovered based on the expected values generated by perturbing the network.

**Figure 5.**
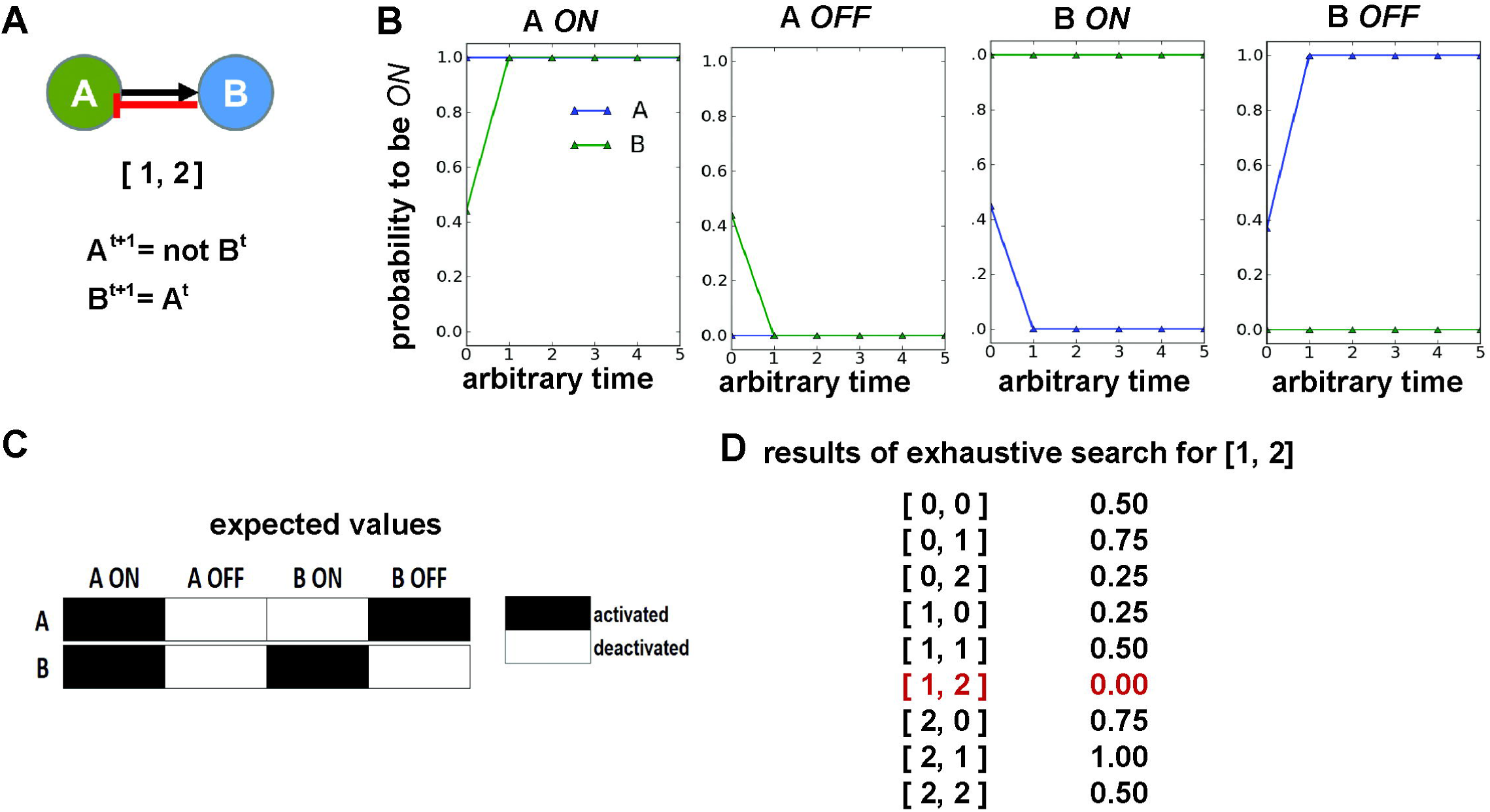
Results of the **[1,2]** network search. **(A)** Network **[1, 2]** is made of the following logic regulations: **A^t+1^ = not B^t^ and B^t+1^ = A^t^. (B)** The expected values of network **[1, 2]** were generated based on the steady state values of nodes A and B under the following perturbation tests: (1) **A *ON***, (2) **A *OFF***, (3) **B *ON***, and (4) **B *OFF***. **(C)** In this search, the p-score of each of the 9 logic networks was calculated from the simulated output of each network and the expected values of network **[1, 2]**. As expected, in this example only network **[1, 2]** had a p-score of 0.0, indicating an exact match.

### Three-node test network

The addition of a single node increases the size of the potential search space considerably. In a 3-node circuit consisting of **A, B**, and **C** (**Figure 4B**), there are 729 interactions and 2,197 logical networks possible (**Table 1**; and **Supplemental Table 2**). Expected values were generated *in silico* for 3 test networks: **[9, 1, 0]**, **[1, 3, 11]**, and **[3, 0, 10]**, where each network is encoded as **[A_i_, B_i_, C_i_]**, and *i* indicates a unique logic rule identifier possible for a given node (**Figure 4B**). Searches for the 3 three-node networks were run using both a GA and an exhaustive search (where all possible 2,197 logic networks were simulated). In all cases, a p-score was calculated for each candidate network tested. Any network with a p-score of 0.0 was reported as a possible network solution.

Three separate sets of *in silico* perturbation experiments (**Table 5**) were used to generate expected values for the three-node networks tested. The correct network was recovered from the expected values of all 3 sets of *in silico* perturbation tests for all 3 test network searches. Perturbation test sets I and II, however, produced non-unique solutions (with many other networks producing a p-score of 0.0). In contrast, perturbation test set III uniquely identified the correct network in all 3 target network searches (**Table 5**).

**Table 5.** Perturbation tests used to search for three-node networks. Results are summarized for three network searches: **[9, 1, 0], [1, 3, 11], and [3, 0, 10]**. Networks are encoded as **[A_i_, B_i_, C_i_]** where **i** indicates the regulation rule present in the network for each node (see **Supplemental Table 2** for the list of regulation rules by node).

| Test Set | Perturbation tests in set | correct network found? / unique solution found? |  |  |
| --- | --- | --- | --- | --- |
|  |  | [9,1,0] | [1,3,11] | [3,0,10] |
| I | 1. **B OFF** | Yes / No | Yes / No | Yes / No |
| II | 1. **A OFF and B ON**<br>2. **A OFF and C ON**<br>3. **A ON and B OFF**<br>4. **A ON and C OFF** | Yes / No | Yes / No | Yes / No |
| III | 1. **A ON**<br>2. **A OFF**<br>3. **B ON**<br>4. **B OFF**<br>5. **C ON**<br>6. **C OFF** | Yes / Yes | Yes / Yes | Yes / Yes |

The three-node network searches performed via a GA were initialized with a population size of 100 and were allowed to run for 20 generations, requiring up to 2,000 simulations for each network search. While 2,000 is only slightly smaller than the 2,197 simulations needed to perform an exhaustive search, the number of simulations performed in 20 generations by the GA was always less than 2,000 because of optimizations in the algorithm that prevent unnecessary repeated simulations of candidate networks maintained in multiple generations of the GA. In all cases, the GA was more efficient at recovering the target network (**Table 6**). Using perturbation test set III, the correct network was found on average in 6 or fewer generations (**Table 6**). The longest GA search (for target network **[3, 0, 10]**) took an average of 41.7 minutes to complete 20 generations, while an exhaustive exploration of the full search space took an average of 98 minutes, under otherwise identical computing conditions.

**Table 6.** Comparison of the time to find the correct network in three different three-node searches. Each search was run three times via genetic algorithm (GA). In addition, exhaustive evaluation of all 2,197 networks in the search space was performed three times. All simulations were distributed across 5 Nehalem core processors. GA tests were initialized with population size of 100 and ran for 20 generations. Values listed are the mean ± the standard deviation of 3 replicates of each network search. All searches used perturbation test set III, which is summarized in Table 5.

|  | Search for<br>[9, 1, 0] | Search for [1,<br>3, 11] | Search for [3,<br>0, 10] |
| --- | --- | --- | --- |
| generation target network<br>found in GA search<br>(mean ± std) | 3.0 ± 2.0 | 4.3 ± 2.5 | 5.7 ± 4.0 |
| elapsed minutes to run 20<br>generations in the GA search<br>(mean ± std) | 40.0 ± 12.5 | 36.3 ± 10.0 | 41.7 ± 9.2 |
| elapsed minutes to run<br>exhaustive search<br>(mean ± std) | | 98.0 ± 26.9 | |

### Additional test networks

We performed similar tests using increasingly complex networks. Refer to the supplemental material for a detailed discussion about how we searched for a predefined Five and Ten Node network (**Supplemental Figure 3, Supplemental Figure 4**, respectively).

### Unraveling the MCF10A network

Returning to the original hypothesized network of MCF10A signaling, we performed additional western blot experiments to obtain readouts representative of mammalian target of rapamycin complex 1 (mTORc1) signaling, which may play a fundamental role in the cross talk between the MAPK and PI3K signaling pathways (**Figure 2A-B**) (Foster and Fingar 2010; Hay 2005; Sarbassov et al. 2006; Sarbassov et al. 2005). Specifically, we measured the phosphorylation of raptor, mTOR and TSC2 under the same six experimental conditions used previously with MCF10A cells (**Figure 6A**). From these data, we concluded that mTORc1 is activated in all 6 experimental conditions, while the TSC1/2 dimer is inactive in all 6 experimental conditions. The activation of mTORc1 was inferred ***ON*** in all 6 conditions because a band was observed in all conditions for phospho-mTOR and phospho-raptor. While the band for phospho-mTOR was weakest in the control condition, we assume that its activity is above the threshold needed for functional activity. For simplicity, we also assumed that the other key molecular components of mTORc1 are present and in an active conformation. The activation of TSC1/2 was inferred ***OFF*** in all 6 conditions because a band representing phosphorylation to TSC2 at T1462 was observed in all 6 experimental conditions. Phosphorylation of TSC2 at T1462, which inactivates the molecule, is regulated by AKT. This residue was phosphorylated even in conditions where AKT was not activated, however. We therefore hypothesized that TSC2 is either constitutively deactivated in MCF10A cells or another key EGFR driven regulator of this residue is missing from our model.

**Figure 6.**
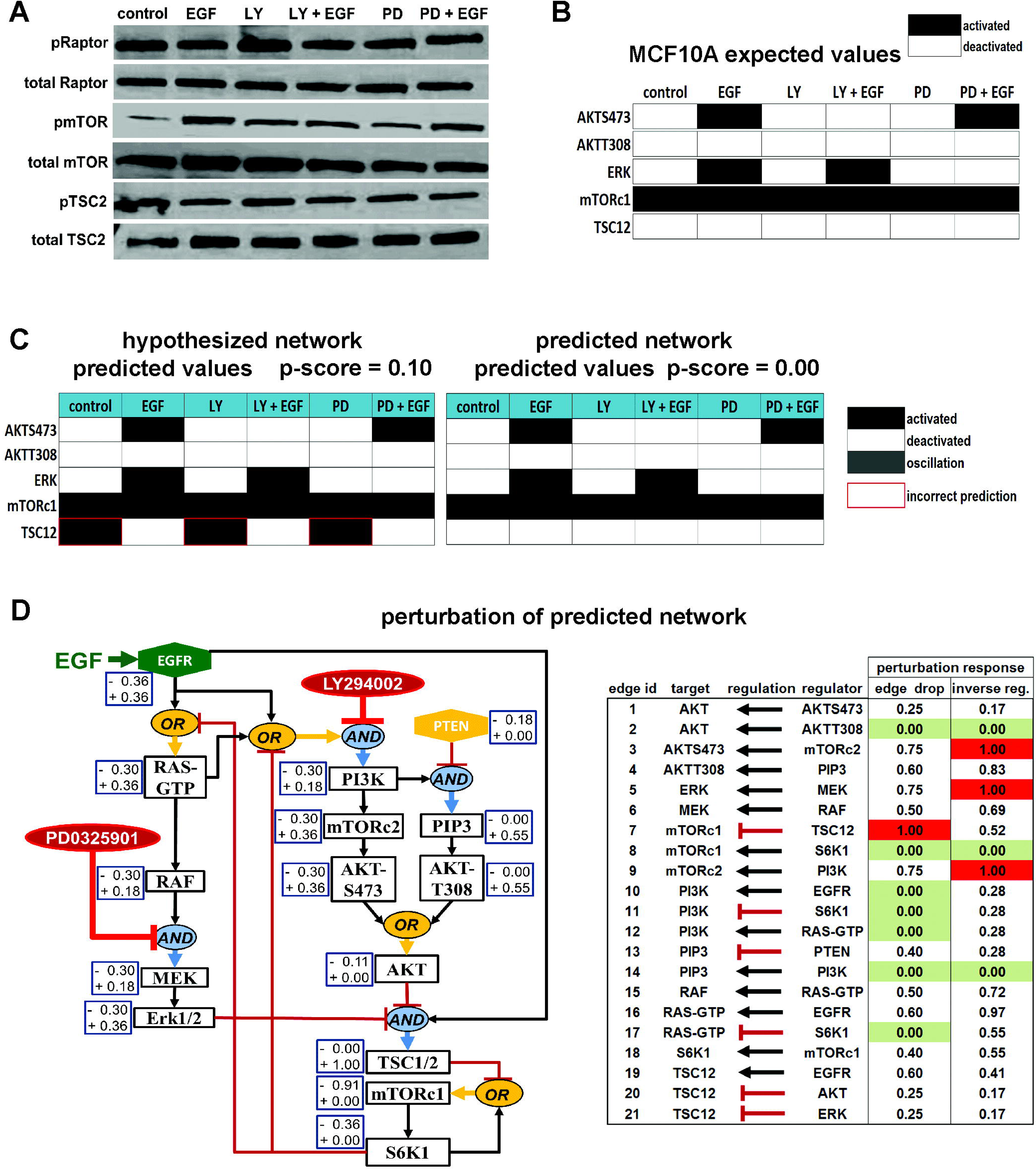
Predicted network of epidermal growth factor receptor signaling in MCF10A cells. **(A)** Western blot of total and phosphorylated Raptor, mTOR, and TSC2 in MCF10A cells in six experimental perturbation conditions are presented along with **(B)** the discretized form of these data as logical expected values. Expected values for the **AKTS473, AKTT308**, and **ERK** network nodes are also included for completeness. **(C)** Based on literature knowledge and a series of constrained genetic algorithm (GA) searches, we predict that the network in Figure 2B should also include an EGFR driven activator of TSC12. The predicted network generated a p-score of 0.0. **(D)** A perturbation analysis of the predicted network was performed. A perturbation response (the p-score after a single network perturbation divided by the maximum p-score produced by all perturbations of the same type) was calculated for each perturbation. The results of a node-based perturbation analysis (where a single node is set to constitutively active and then constitutively inactive) is summarized. The boxes next to nodes list the perturbation response when the node was perturbed *OFF* (-) and *ON* (+). The table on the right summarizes the results of two different perturbation analyses: an edge-based perturbation analysis where one regulatory edge is removed and a regulatory-based perturbation analysis where each edge is inverted to its opposite regulatory type (e.g., activation to inhibition and inhibition to activation). Perturbations that had no effect on the network’s p-score are highlighted in green (0.00), and perturbations that had the greatest effect on the network’s p-score are highlighted in red (1.00).Tables

When the expected values for **mTORc1** and **TSC12** activation were added to the expected values for **AKTS473**, **AKTT380**, and **ERK** (**Figure 6C**), the p-score of the original MCF10A hypothesized logic network (Figure 2B) increased from 0.00 to 0.10. The change in p-score reflects the difference between the simulated values of the hypothesized network and the expected value for **TSC12** activation in the control, LY, and PD conditions (**Figure 6C**). We used the GA to search for networks that are consistent with the MCF10A expected values in **Figure 6C.** First, we used the GA (generations=100, population=400) to search the informed search space described previously as consisting of 275 distinct logic rules and **2.4 x 10^12^** distinct logic networks (**Supplemental Table 1**). While more than 1,000 networks with a p-score of 0.0 were found, they all included one of three regulation rules for TSC1/2, each of which violated our literature derived knowledge of AKT and/or ERK inhibition of TSC1/2 (Hahn-Windgassen et al. 2005; Hay 2005; Jiang and Liu 2008; Manning et al. 2002; Sabatini 2006). The three TSC1/2 rules contained in all networks found with a p-score of 0.0 were: (1) **TSC12^t+1^ = AKT^t^**, (2) **TSC12^t+1^ = AKT^t^ and not ERK^t^**, and (3) **TSC12^t+1^ = AKT^t^ and ERK^t^**.

We next searched a larger search space of **5.7 x 10^14^** possible networks where **EGFR** was added as a potential direct regulator of **mTORc1**, **S6K1**, and **TSC12**. The inclusion of **EGFR** as a possible regulator of these three nodes allowed us to test for the presence of molecules that regulate each of these nodes as a direct consequence of **EGFR** activation without including additional nodes in the network. In this search, more than 5000 networks with a p-score of 0.0 were found in the first GA run (suggesting there are likely tens of thousands of networks in this search space that produce steady state values that match the expected values). If we could measure the activation of every protein in the network under all six perturbation conditions, the number of networks consistent with our data would likely be far smaller than the very large set of networks found with a p-score of 0.0. To attempt to reduce the set of candidate networks in this large search space without performing additional experiments, constraints were added to the GA for node regulations that we had high confidence in, based on the literature and our experiments. Specifically, the constraints required the following to be true:

1. ERK was only activated by **ERK^t+1^ = MEK^t^**,
2. RAF was only activated by **RAF^t+1^ = RAS-GTP^t^**,
3. PIP3 was only activated by **PIP3^t+1^ = PI3K^t^ and not PTEN^t^**.

In addition, rules that met any of the following criteria were removed from the search space because they violate what we expect in normal canonical signaling in these pathways (**Table 3**):

1. any potential regulation of **mTORc2** that included inhibition by **PI3K**.
2. any potential regulation of **AKT** that included inhibition by the phosphorylation of its T308 or S473 residues.
3. any potential regulation of **RAS-GTP** that included inhibition by **EGFR** or activation by **S6K1**.
4. any potential regulation of **PI3K** that included inhibition by **EGFR** or inhibition by **RAS-GTP**.

While the inclusion of these constraints reduced the effective search space by 5 orders of magnitude to 2.8 X 10^9^, this still represents an extremely large search space. When we searched under these constraints, thousands of networks were again found with a p-score of 0.0. One of the networks found, however, was identical to the original hypothesized network of normal signaling (**Figure 2B**) in all but the **TSC12** regulation. The **TSC12** regulation in this network was **TSC12^t+1^ = EGFR^t^ and (not AKT ^t^ and not ERK^t^)**. The presence of this network is significant because it does not violate our literature knowledge of TSC1/2 inhibition by AKT and/or ERK, and it suggests that if the rest of our hypothesized network regulations are correct in **Figure 2B**, then an EGFR driven molecule other than AKT and ERK is also important for the regulation of TSC1/2 activation in MCF10A cells. Because of the closeness of this network to our original hypothesized network, we performed a perturbation analysis on this network (**Figure 6D**, **Supplemental Figure 5**). In the perturbation analysis, a single perturbation is made to the logic network in the form of an edge deletion or an inverse regulation, and the resulting is p-score calculated. The reported perturbation response is calculated by dividing the p-score of the perturbed network by the maximum p-score found by all perturbations of a given type (e.g., node, logic operator, edge, or edge regulatory type). The most important regulations in the network were related to the regulation of mTORc1 and TSC1/2 as well as MEK activation by ERK and mTORc2 activation by PI3K.

Finally, we found that if additional constraints were added to the GA that violated literature knowledge of signaling interactions in this network, then we were ultimately able to reduce the number of networks with a p-score of 0.0 to a set that numbered in the hundreds rather than the thousands. A large number of these networks are the products of experimental uncertainty in some interactions in the network. For example, because T308 is never phosphorylated on AKT in MCF10A cells (almost certainly because of the presence of activated PTEN in these cells) (**Figure 2C**), without additional experiments that directly perturb this residue in some way, our methodology is not able to distinguish between the following two regulations for AKT activation:

> **AKT^t+1^ = AKTS473^t^**
>
> **AKT^t+1^ = AKTS473^t^ or AKTT308^t^**

Similarly, because PTEN was activated in all conditions in our experiments, the presence of active PI3K is logically unimportant (because inhibitory **PTEN** dominates activating **PI3K** in one of the rules), and thus the methodology cannot distinguish between these two regulations for PIP3:

> **PIP3^t+1^ = not PTEN^t^**
>
> **PIP3^t+1^ = PI3K^t^ and not PTEN^t^**

Moreover, the uncertainty in these regulations is reflected in the 0.0 perturbation response when the **AKTT308** to **AKT** or the **PI3K** to **PIP3** edge was dropped in the perturbation analysis (**Figure 6D**).

## Discussion

When testing that a network inference approach is predictive, it is important to test it in a controlled a way. We inferred the EGF-driven PI3K and MAPK network circuity inside MCF10A mammary epithelial cells, and then evaluated the accuracy of our predicted network with in silico tests against a target model network that was known a priori within live cells. In the case of MCF10A cells, we relied on phospho-western blot data generated from a series of perturbation experiments performed with and without EGF stimulation as the input (in the form of expected values) to the search algorithm.

Our results indicate that the method can be used to help infer the underlying signaling circuitry in a population of cells if sufficient experimental data are available. The approach developed can be readily applied to very large networks, but to perform a series of GA searches for a large network, access to high-performance computing facilities is recommended. A limitation of our approach is the implementation of a heuristic GA algorithm, which does not exhaustively search the network space. While synchronous simulation takes considerably less time to run than the asynchronous simulation approach we used, our testing revealed that synchronous simulations could not always find the expected network in a search space. For *in silico* tests, we generated expected values from known test networks and used these values as input to the search algorithm. The tests suggest that asynchronous logic simulations and a GA search can theoretically be used to identify an unknown network from the output of a set of network perturbations. Future development efforts are planned to further optimize the algorithm and ideally reduce the computing time necessary to converge to an optimal solution in large search spaces.

Our results also suggest that the more experimental perturbations performed, the more likely discovery of the source network is. We previously illustrated this point with a systematic perturbation analysis to infer the effects of Honokiol on the Notch signaling pathway in SW480 colon cancer cells (Wynn et al. 2014). Not surprisingly, we also observed that it was almost always necessary to obtain a readout of all non-input nodes in the network in order to converge on a single correct network in a search space. In a perturbation experiment, the state of all input nodes should be known *a priori*. Otherwise, it is not appropriate to treat the node as an input node. It is also clear that as the number of nodes in a network increases, the possible ways the nodes may be connected in a logic network becomes extremely large. Therefore, if attempting to infer the signaling network responsible for observed experimental readouts in a biological network, an exhaustive exploration of the search space of all possible logical circuits will almost never be possible. To overcome this challenge, our results strongly suggest that a GA can provide a reasonable heuristic-based alternative to an exhaustive search. It should be emphasized, however, that even with the search optimization provided by the GA, to obtain results in less than a day for large search spaces (such as the ones used for the 10 node test network or the 15 node hypothesized MCF10A network), it was often necessary to distribute the simulations across up to 40 computing cores.

We relied on phospho-western blots for protein activation readouts. An alternative high-throughput approach for measuring protein activation is the use of multiplexed bead-based immunoassays, which consist of a set of antibodies bound to beads with individual fluorescent signatures, allowing detection of phosphorylation sites (Du et al. 2009; Krishhan et al. 2009). Because the multiplex system can detect several analytes in a single sample volume in far less time than equivalent data can be collected from traditional western blot experiments, this approach holds tremendous potential when used in tandem with protein network inference approaches. Saez-Rodriguez et al. (2009) used the continuous median fluorescence intensity readout of phospho bead-based immuno assays as input to another discrete Boolean based network inference approach. While their method differed from ours in a few important ways, the most significant difference is their use of synchronous Boolean simulations. It is worth noting that their inference approach also used a GA, providing additional support for the utility of heuristic based approaches for limiting the enormous network search spaces encountered in these types of problems.

Our approach recovered the target model network in all *in silico* tests performed. In some cases, it was necessary to perform additional *in silico* perturbation experiments in order to identify a single unique and correct network with a p-score of 0.00. In the case of the 5 node network, the target model network could only be distinguished in the informed search space. When we used no prior knowledge of the regulation of **B, C, D, or E** (the uninformed search space), a set of 52 networks with the same or nearly the same steady values under all possible perturbations combinations was found (**Supplemental Figure 3**). If this situation is encountered with a real network in the laboratory, we recommend to expand the initial hypothetical network with additional nodes, which are considered critical to the signaling mechanism.

Similarly to Saez-Rodriguez et al. (2009), we did not identify one single logic network that matched our experimentally measured data. To do so would require performing many additional experimental perturbations as well as obtaining readouts of all or most activation states of the proteins in our hypothesized network, which was based on an extensive literature search. It is noteworthy that we did find a single network that matched the experimental data for all conditions tested, except for the logical regulation of TSC1/2. Our hypothesized network originally assumed that other regulators of TSC1/2 were unimportant relative to AKT and ERK. The final predicted network (**Figure 6D**) suggests that this is not the case and that another EGFR-regulated molecule may play an important role in TSC1/2 regulation. While this result stops short of inferring the most likely “target model” network inside MCF10A cells, it provides a tangible prediction for additional laboratory explorations related to the regulation of TSC1/2.

Of course, finding the “target model” network inside a living cell may not be fully attainable for any reverse engineering methodology and it may not even be possible, given the heterogeneity expected within a population of cells. In reality, what we are likely to find are mechanisms and models of a set of integrated pathways that have not yet been contradicted by experiment. This will be true of all modeling frameworks, not just Boolean based network models. Boolean logic, however, provides a predictive approximation of the transfer of signals in a circuit and has been used to build predictive models of the cell cycle (Davidich and Bornholdt 2008; Li et al. 2004; Ribba et al. 2006), gene regulatory networks (Albert and Othmer 2003; Chaves et al. 2005), circadian clocks (Akman et al. 2012), and signaling networks (Li et al. 2006; Zhang et al. 2008). Living cells behave as extremely robust integrated circuits that exhibit significant heterogeneity, but still respond to external stimuli in predictable ways. The deterministic cellular outcome, for example the induction of proliferative programs in response to growth factor stimuli or the induction of p53 oscillations in the presence of DNA damage, has been compared to digital biological computation (Lahav et al. 2004; Tamsir et al. 2011).

The approach presented here provides a means for inferring and/or validating a network hypothesis from experimental data. Our approach could be used to identify novel adaptive resistance mechanisms of cancer cells to therapeutic drug targets. The reverse engineering methodology presented here can also be implemented to investigate the potential differences between normal and cancer cell lines. Ultimately a logic-based network model can provide only qualitative dynamics of the predicted order of signaling events but cannot, as it stands, provide precision in terms of timescales or the size of a response. Two-state Boolean models can predict that a molecule is activated but not the degree of activation or signal amplification. While more precise approaches with respect to time and signal amplitude are widely used in systems biology, the predictive power of these methods is substantially limited by their dependence on a large number of unknown parameters and kinetic mechanisms (Mourao et al. 2011; Srividhya et al. 2007; Wang et al. 2013; Wynn et al. 2012) that must be assumed prior to simulation. Thus, despite some limitations, when judged in the context of other methods, logic network models provide a predictive and robust methodology for modeling biochemical regulation without requiring prior knowledge of rate constants and reaction mechanisms.

## Acknowledgements

SS and MLW acknowledge support from the James S. McDonnell Foundation’s grant support for studying complex systems. This work was also partially supported by NIH R25 DK088752 (NC and SS), the University of Michigan Protein Folding Diseases Initiative (SS), the Breast Cancer Research Foundation (MLW, ME, ZFW, and SDM), and the Avon Foundation (ZFW and SDM).

